# Dopamine enhanced auditory perceptual learning in humans via long-term memory consolidation

**DOI:** 10.1101/2021.06.16.448678

**Authors:** Ding-lan Tang, Jun-Yun Zhang, Xiaoli Li, Yu-Xuan Zhang

**Author notes:** **Author Contributions** D-L.T. and Y-X. Z. designed the experiments, analyzed the data and prepared the manuscript. D-L. T. conducted the experiments. All authors edited the manuscript for publication.

## Abstract

Dopamine is known to modulate sensory plasticity in animal brain, but how it impacts perceptual learning in humans remains largely unknown. In a placebo-controlled, double-blinded training experiment with young healthy adults (both male and female), oral administration of Madopar, a dopamine precursor, during each of multiple training sessions was shown to enhance auditory perceptual learning, particularly in late training sessions. Madopar also enhanced learning and transfer to working memory when tested outside the time widow of drug effect, which appeared to retain for at least 20 days. To test whether such learning modulation was mediated by the dopaminergic working memory network, the same dopamine manipulation was applied to working memory training, but to little influence on learning or transfer. Further, a neural network model of auditory perceptual learning revealed distinctive behavioural modulation patterns for proposed dopaminergic functions in the auditory cortex: trial-by-trial reinforcement signals (reward/reward prediction error and expected reward) and across-session memory consolidation. Only the memory consolidation simulations matched experimental observations. The results thus demonstrate that dopamine modulates human perceptual learning, mostly likely via enhancing memory consolidation over extended time scales.

## Introduction

Dopamine, a neurotransmitter widely present in the brain, has long been known to play critical roles in neural computations underlying learning and behavioral adaptations ^1–5^. One of the most remarkable demonstrations of dopamine’s function in modulating neural plasticity is in the auditory cortex, where electrical stimulation of midbrain dopamine neurons coupled with pure tones induced stimulus-specific changes in neuronal responses and reorganization of the tonotopic map in the rat brain ^6^. Follow-up animal studies link the dopaminergic influence on the auditory cortex to auditory perceptual learning and memory functions ^7–10^. Note that perceptual learning and memory formation in animals rely on instrumental conditioning, in which sensory stimuli are associated with an appetitive or aversive signal. Perceptual learning in humans, however, occurs with sensory experiences without apparent conditioning signal, even for background stimuli presented below the perceptual threshold ^11^ or in the absence of sensory difference to discriminate^12^. However, the dopamine effect on auditory perceptual learning and memory in humans remains to be examined.

Dopamine precursor drugs such as levodopa have been used to examine perceptual, cognitive and motor consequences of dopamine in the human brain. For example, oral administration of levodopa, which elevates plasma dopamine level on the time scales of minutes to hours, has been shown to enhance incidental learning of motor sequence in patients with Parkinson’s disease, motor memory formation in healthy and aging individuals ^13,14^, and associative learning of spoken words in healthy adults ^15,16^. Levodopa was also reported in a functional magnetic resonance imaging (fMRI) study to enhance BOLD activity in the left auditory cortex during reward learning, but without behavioural significance ^17^. These studies typically involve a single administration of dopamine drugs, while perceptual learning often takes place across multiple training sessions spanning over days or weeks ^18,19^.

There is more than one possible mechanism for dopamine drugs to affect perceptual learning. The most influential one is possibly reward prediction error (RPE) encoded by subsecond activities of midbrain dopamine neurons, namely discrepancy between expected and actual reward, serving as a reinforcing signal for organisms to learn by trial and error ^20–23^. Such learning has hence gained the name “reinforcement learning”. Further, dopamine changes may impact learning via their influence on cognitive functions such as working memory that involve the prefrontal and striatal dopaminergic networks ^4,5^. Supporting this view, human working memory performance can be modulated by dopaminergic drugs in a dose dependent manner ^24–26^. Patients of Parkinson’s disease with striatal dopamine deficits showed impairment in a form of reward learning that correlated with working memory performance and the impairment could be remediated by dopaminergic medication ^27^. Last but not the least, dopamine has been shown in gerbils undertaking multiple-session training on frequency modulation discrimination to affect only the late sessions of learning via protein synthesis dependent mechanisms, presumably memory consolidation ^7^.

Here, we examined how dopamine drugs administered during multi-session training affected auditory perceptual learning, in two separate, placebo-controlled, double-blind training experiments with young, healthy participants combined with neural-network modeling of RPE-weighted reinforcement learning. Madopar (100 mg of dopamine precursor levodopa plus 25 mg benserazide, a dopamine decarboxylase inhibitor) was orally administered before each of 7 to 9 daily training sessions. Learning and transfer were evaluated at least 24 hours before and after drug administration to be free of temporary effects of plasma dopamine elevation. In the first experiment, dopamine effect was tested on auditory perceptual learning using a tone frequency discrimination task, for which we have previously shown learning over multiple daily sessions of repeated practice and bi-directional transfer with working memory learning ^28,29^. In the second experiment, dopamine effect on working memory learning was examined using an auditory version of the n-back task that has been reported to demonstrate dopamine-dependent learning ^26,30^. A neural network model employing RPE-weighted reinforcement learning for frequency discrimination was constructed to simulate drug effects on learning and consolidation mechanisms. The experimental results indicate that dopamine drug can enhance multiple-session auditory perceptual learning as well as its transfer to working memory, but the same drug manipulation fails to enhance working memory learning *per se*. Modeling simulations reveal that the observed dopaminergic modulation of auditory learning is unlikely to result from RPE-related phasic dopamine effects, but similar to enhancement of across-session memory consolidation.

## Materials and methods

### Participants

Forty-four healthy young participants (23.55 ± 4.82 years, 40 females) volunteered for a double-blind, placebo-controlled study with dopaminergic drug, approved by Peking University Institutional Review Board. All participants had normal hearing (⇐20 dB HL bilaterally on pure tone audiometry between 0.5 to 4 kHz) and reported no history of neurological, psychiatric, or cardiological disorders, chronic or acute diseases, or use of drugs affecting the central nervous system up to 2 weeks before participating in the study. Participants signed consent forms and received compensation for their participation.

The participants were randomly assigned to either a dopamine or a placebo group for one of two multi-session training experiments. Those in the dopamine groups orally took 125 mg Madopar (100 mg Levodopa plus 25 mg benserazide) before each training session, while those in the placebo groups took vitamin with similar volume and appearance. Group identity was assigned and drug/placebo were prepared by a third party, keeping both the participants and the experimenter blind. One participant withdrew before completing the training experiment for personal reasons, leaving a total of 19 participants in auditory learning Experiment (10 in the dopamine group, 9 in the placebo group) and 24 participants in WM learning Experiment (12 in the dopamine group and 12 in the placebo group).

After drug administration, participants’ vital signs including heart rate and skin conductance were monitored during the experimental session. Participants were instructed to stop participation and receive medical care from a residing internal medicine specialist if they felt nausea, headache, or any other possible drug side effects, or displayed abrupt changes in their vital signs. After training, the participants completed a questionnaire regarding their well-beings. None of the participants reported symptoms serious enough to interrupt the experiment.

### Experimental design

Effect of dopaminergic drug on learning was examined in two multi-session training experiments: one for auditory discrimination training and one for working memory training. Each experiment consisted of a pre-test, multiple training sessions (7 for auditory discrimination training and 9 for working memory training) and a post-test on consecutive days (Fig. 1). Approximately half of the participants returned for a retention test 20 days after the post-test. In the pre-, post- and retention tests, performance on the training and transfer tasks was assessed (details of the tasks are described in Stimuli and Task). Drug/placebo were only administered in training sessions. The amount of Madopar (100 mg Levodopa and 25 mg benserazide) was chosen as the lowest among previous studies using similar drugs with healthy adults ^15,31–33^, which produces peak plasma level of dopamine approximately 1 hour after intake, with an elimination half time of 90 minutes. To maximize increase of dopamine level during training, each training session was truncated to approximately 30 minutes and started 40 minutes after oral intake. Pre- and posttests were separated from drug administration by at least 24 hours and thus should be free of drug influence.

**Figure 1.**
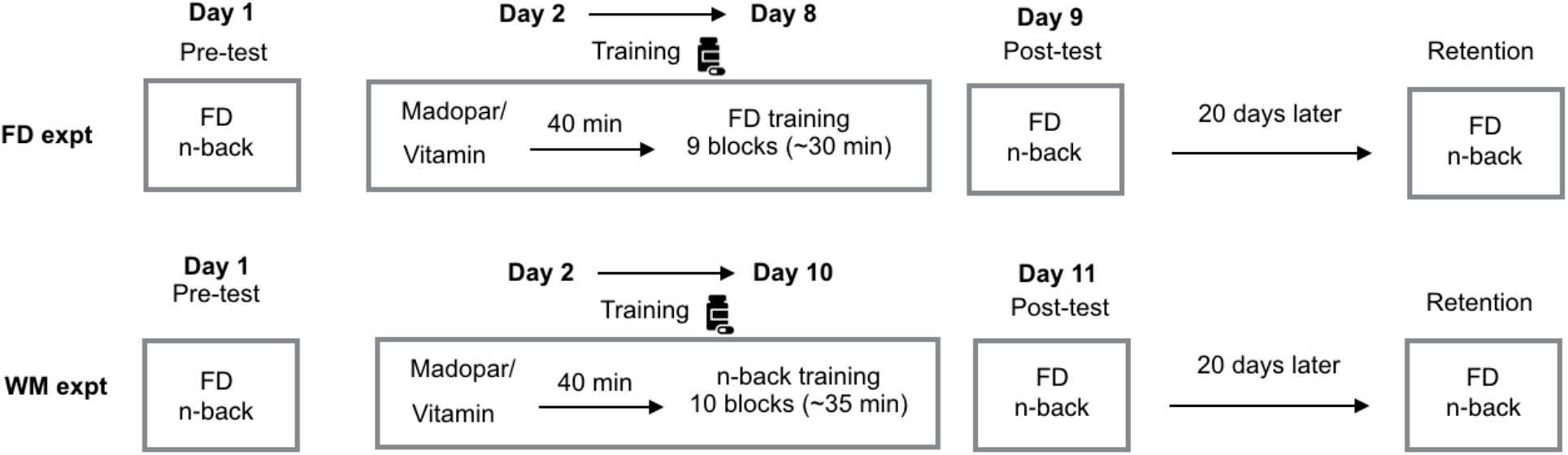
Experimental design. A pretest-training-posttest-retention design was used for both auditory discrimination training and working memory training experiments. FD: tone frequency discrimination; WM: working memory; n-back: Tone n-back.

The training procedures were adopted from a previous study demonstrating learning with the same training tasks ^28^. For auditory discrimination training, participants practiced on Tone Frequency Discrimination (FD) for 9 blocks of 50 trials (approximately 30 minutes) in each of seven training sessions. For working memory training, participants practiced on Tone n-back for 10 blocks of 40+n trials (approximately 35 minutes) in each of nine training sessions, with n starting at 2 and progressing to 3 after 2-back performance either improved for 3 consecutive sessions or exceeded 90% correct for 5 out of 8 consecutive blocks. The FD and n-back tasks were assessed in all test sessions for both experiments, each serving as training task for one experiment and test task of learning transfer for the other experiment.

### Task and Stimuli

#### Tone frequency discrimination (FD) task

Auditory discrimination was measured and trained with a tone frequency discrimination (FD) task ^28^. Both testing and training were conducted in blocks of 50 trials, with each block yielding a measurement of discrimination threshold. In each trial, three tones were presented sequentially in a random order with an inter-stimulus interval of 300 ms, two of which were identical (standards) and one had a higher frequency (the target). Participants were instructed to indicate the position of the target tone by pressing a button on the computer keyboard. Trial-by-trial feedback was provided visually during training, but not in testing sessions. All tones were 100 ms long, including 15-ms rise/fall ramps, presented diotically at 75 dB SPL via Sennheiser HD-499 headphones (Sennheiser electronic GmbH & Co. KG, Wedemark, Germany). The standard frequency varied randomly from trial to trial between 0.9 and 1.1 kHz in 50 Hz steps.

In each block, the starting frequency difference between the target and standard tones (ΔF, expressed as percentage of the standard frequency) was 50%, and was divided (for a correct response) or multiplied (for an incorrect response) by a factor of 2 until the first reversal. After that, ∆F was varied following a 3-down 1-up rule, with a step size of 1.41, to estimate FD threshold at 79% correct point on the psychometric function ^34^. A demo of 5 trials was used to familiarize the participants with the task before the first run.

#### Tone n-back

Auditory working memory was measured using an n-back task with tonal stimuli ^28^. In each test session, 2 blocks of tone 2-back and 2 blocks of tone 3-back were administered. In training, n started with 2 and progressed to 3 according to the criterion aforementioned. For each block, a sequence of 40 + n tones (43 tones for 3-back and 42 tones for 2-back) was presented with an ISI of 2,500 ms. Participants were instructed to press a button only if the current tone matched the tone from n steps earlier in the sequence (a target). Each sequence contained 12 targets randomly distributed in the last 40 tones. All tones were presented diotically at 60 dB SPL. Tones included in each sequence were of eight frequencies drawn between 1,080 to 4,022 Hz and separated by at least one equivalent rectangular bandwidth, so that they can be easily distinguished by participants. During training, visual feedback was provided after each button press, and performance in percent correct was displayed at the end of each sequence. A demo of 20+n trials was used to familiarize the participants with the task before the first run of 2 and 3 back tasks.

### Neural network model of FD learning

The FD learning model (Fig. 2) consists of three layers: a sensory layer, a memory layer, and a decision-making layer (single unit). The sensory layer is a set of units that respond differentially to sounds with different frequencies, forming a tonotopic map as observed in the primary auditory cortex ^35,36^. Firing rate of sensory unit i for an input tonal stimulus with frequency fs is given by

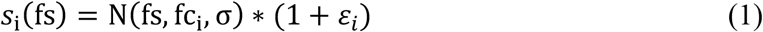

**Figure 2.**
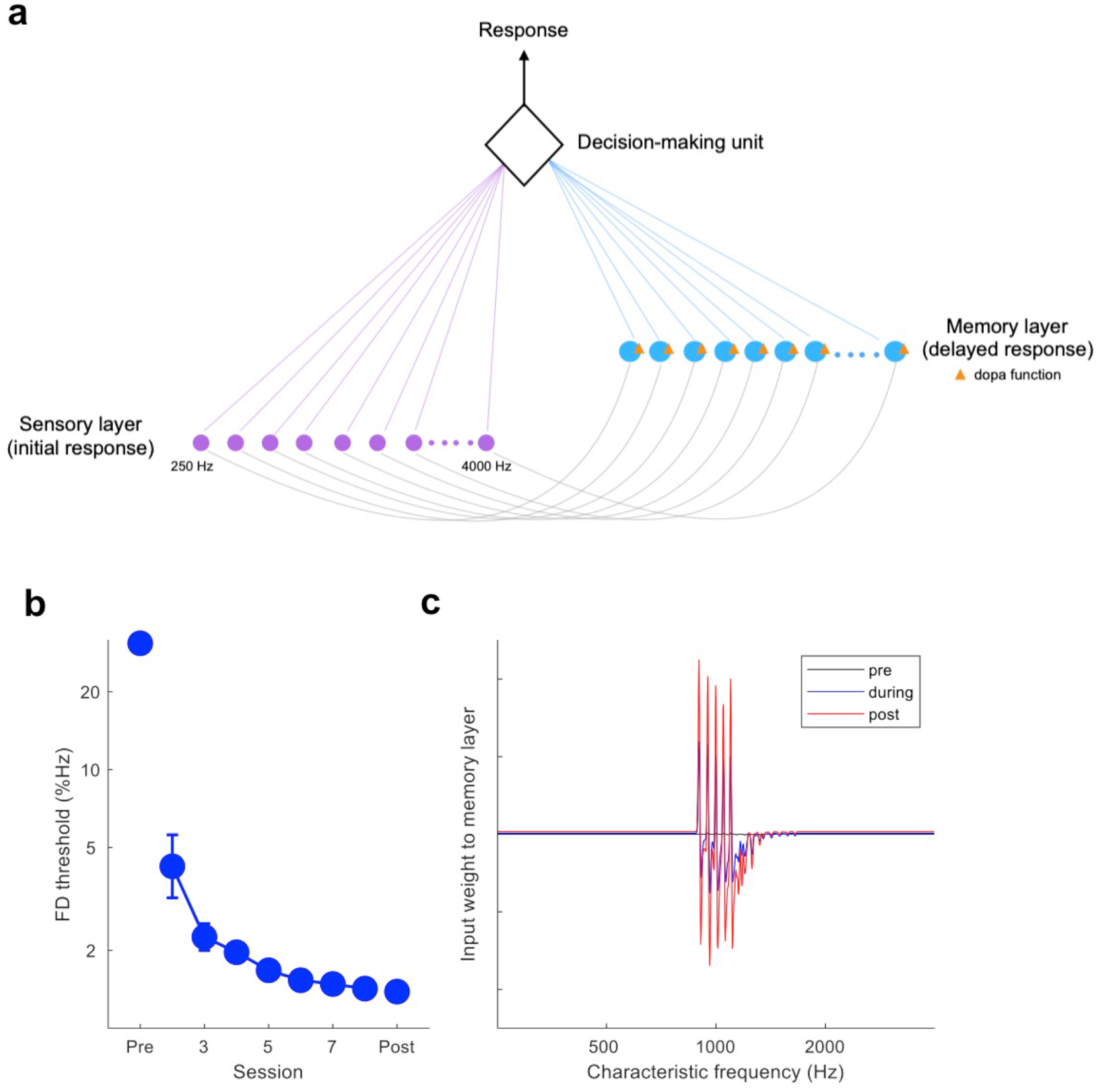
Neural network model of FD learning. (a) Illustration of the model structure. FD discrimination of two sequentially presented sounds were made by the decision-making unit by comparing the input from the sensory layer (initial response) for the second sound with that from the memory layer (delayed response) for the first sound. Reward prediction error (RPE) based reinforcement learning takes place in the input weight from the sensory layer to the memory layer, where dopamine functions apply. (b) Baseline learning of the model. FD discrimination by the model started from chance and improved with trial-by-trial feedback, approaching human level at the first training session. Parameters used to generate baseline learning were fixed in subsequent model simulations except for those reflecting dopamine functions. (c) Learning mechanism of the model. Input weight from the sensory to the memory layer starts at a uniform level across the spectrum (arbitrary unit, set at 1 in the current case; black line), generating chance level performance before training, and shows stimulus specific changes during (blue line) and after (red line) training. Specifically, memory input weight is increased at the standard stimulus frequencies, but reduced to a lesser extent at the comparison stimulus frequencies. Such weight changes correspond to enhanced memory for the trained standard stimuli.

Where N is the normal probability density function cantered at the unit’s characteristic frequency fc_i_ with width of σ, and *ε*_*i*_ is the encoding noise. In the current simulations, the sensory layer consisted of 1000 units spanning the spectrum of 250 to 4,000 Hz, i.e., 4 octaves with the training frequency region (around 1 kHz) in the middle. Tuning width σ was set at 2.5 times the difference between two neighbouring characteristic frequencies. Internal noise *ε* was initialized to produce an asymmetric ‘U’ shape of discrimination threshold across the spectrum, similar to those experimentally observed ^37^. The memory layer receives input from the sensory layer and responds according to:

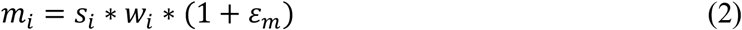

Where *W*_*i*_ is the input weight from the sensory layer to the memory layer, *ε*_*m*_ is the memory noise. The response of the decision unit was given by:

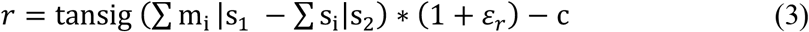

Where *ε*_*r*_ is decision noise, c is the decision criterion. Both *ε*_*r*_ and c were set at 0 for the current simulations, so that decision was made by comparing the summed memory response of the first of two sequentially presented stimuli (s_1_) with the summed sensory response of the second stimulus (s_2_) without bias. The sign of r determines response: positive indicating the first stimulus being higher in frequency (signal), and negative indicating the second being signal. Unlike human listeners, the model starts at chance level of discrimination, and human-like performance can only be obtained with learning.

Learning in this network involves only memory improvement, as the synaptic weight w changes following the Hebbian rule modified by reward prediction error ^38^:

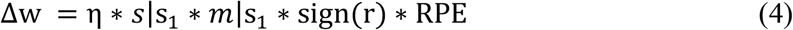

Where η is learning rate, sign(r) is choice of response, RPE is the reward prediction error, calculated as

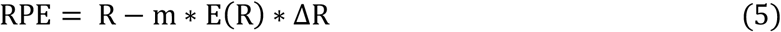

R, the received reward, changes between two levels with feedback: the baseline level of R_0_ for incorrect trials, and increasing by ΔR for correct trials. The coefficient m is set at 1. E(R) is expected probability of reward, given by:

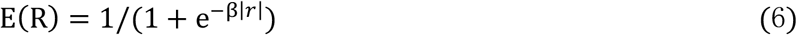

Expected probability of reward is a function of the perceived perceptual difference |r|. The coefficient β represents the propensity of reward expectation. Its value was simulated to generate human-like psychometric functions.

As multiple-session perceptual learning engages across-session or across-day memory consolidation ^39^, the model assumes that learning acquired during a training session would decay following the memory decay function:

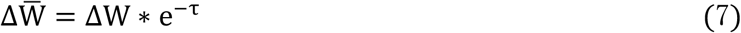

Where ΔW is the weight change (new learning) acquired during a given session,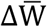 is the weight change retained (consolidated) for the subsequent session. The rate of memory loss, τ, also reflects the level of consolidation: the greater τ is, the less training-induced memory would be consolidated.

### Data Analyses

FD thresholds in percent of standard frequency were log-transformed to approximate normal distribution for data analyses ^28^. Tone n-back performance was measured by d’, calculated as Z(hit rate) – Z(false alarm rate), where Z is the inverse cumulative Gaussian distribution. Statistical analyses were conducted using R (R Core Team, 2019). Learning, transfer, and change in FD-WM relationship were analysed separately using linear mixed-effects models (LME) implemented in the package lme4 ^40^. For learning during training (with drug administration), the model included time (training sessions), group (placebo v.s. Madopar) and time by group interaction as fixed-effects and by-subject random intercepts and slopes as random effects. For learning and transfer in testing sessions (without drug administration), the model included time (pre v.s. post or post v.s. retention), group (placebo v.s. Madopar) and time by group interaction as fixed-effects and by-subject random intercepts as random effects. For FD-WM relationship change, the model included FD performance as dependent variable, WM performance as predictor, WM by test (pre v.s. post) by training (FD v.s. WM trainings) interaction as fixed-effects, and by-subject random intercepts as random effect. Significance of main effects and interactions was decided by using α = 0.05, with t and p values for model terms were obtained by the Satterthwaite’s approximation method implemented in the package lmerTest ^41^. In Results, parameter estimate (est.), standard error (s.e.), t and p values are reported for effects of interests. Post hoc analyses for significant effects were then conducted on the model using the “emmeans” package ^42^.

## Results

### Auditory discrimination learning and transfer were enhanced by dopamine precursor during training

We first examined the effect of Madopar (100 mg L-dopa and 25 mg benserazide), orally administered before each training session, on auditory discrimination performance and learning during those sessions. Both the dopamine and the placebo groups improved significantly on the trained FD task across training sessions (Fig. 3a; Linear mixed-effects model, main effect of time: est. = −0.0763, s.e. = 0.0102, t = −7.45, p < 0.001). Importantly, the dopamine group improved more than the placebo group (Fig. 3a; group by time interaction: est. = 0.0408, s.e. =0.0149, t = 2.74, p = 0.014; main effect of group: est. = −0.0341, s.e. =0.1422, t = −0.24, p = 0.814), demonstrating a learning-enhancing effect of dopamine.

**Figure 3.**
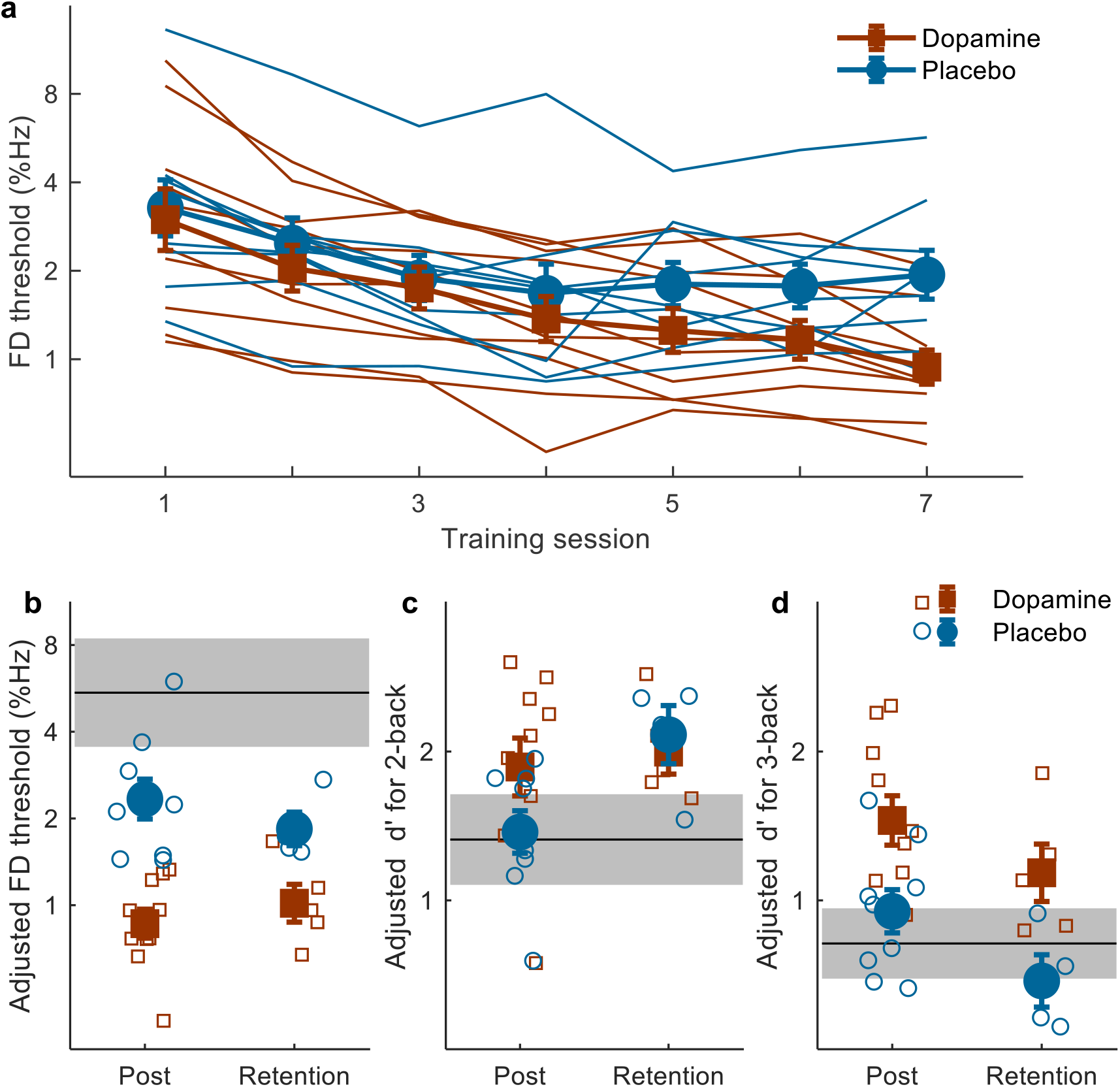
effect of dopamine on FD learning and its transfer to WM. (a) FD performance during 7 days’ training for individuals (thin lines) and group average (thick lines with filled squares/diamonds). Dopamine group (N=10) are shown in orange and placebo group (N=9) are shown in blue. (b, c, d) Individual (open symbols) and group mean (filled symbols) performance for FD (b) and WM 2-back (c) and 3-back (d) performance on post-training and retention (N=5 for dopamine group and N=4 for placebo group) tests, adjusted to account for individual differences in the pretest, with mean pretest performance (black lines) and their 95% confidence interval (grey areas) plotted within each panel. Error bars are ± 1 s.e.m.

Learning and transfer were then evaluated by comparing performance in pre- and posttests, which were separated from drug administration by at least 24 hours and thus were conducted with normal plasma dopamine level. For the trained FD task, both groups improved between the tests (Fig. 3b; main effect of time: est. = 0.8040, s.e. = 0.0992, t = −8.11, p < 0.001), but the dopamine group improved significantly more (group by time interaction: est. = −0.4371, s.e. =0.1441, t = −3.03, p = 0.007; significant main effect of group: est. = 0.4297, s.e. =0.1513, t = 2.84, p = 0.009), indicating enhancement of auditory perceptual learning by dopamine. Post hoc comparisons revealed that there was a significant group difference during posttest (t = −2.84, p = 0.009), but not during pretest (t = 0.05, p = 0.961). For the untrained WM task, both 2- (Fig. 3c; main effect of time: est. = −0.5108, s.e. = 0.1676, t = −3.05, p = 0.007) and 3-back (Fig. 3d; main effect of time: est. = −0.8146, s.e. = 0.1849, t = −4.41, p < 0.001) performance improved significantly between the pre- and posttests. Further, while 2-back improvements were not distinguishable between groups (Fig. 3c; non-significant interaction: est. = 0.4864, s.e. = 0.2435, t = 2.00, p = 0.062; and non-significant main effect of group: est. = −0.1011, s.e. = 0.3310, t = −0.31, p = 0.763), 3-back improvements were greater for the dopamine group than the placebo group (Fig. 3d; group by time interaction: est. = 0.5852, s.e. = 0.2687, t = 2.18, p = 0.044; significant main effect of group: est. = −0.6255, s.e. =0.2344, t = −2.67, p = 0.012). Post hoc comparisons revealed that there was a significant group difference during posttest (t = 2.67, p = 0.012), but not during pretest (t = 0.17, p = 0.864), indicating that elevation of plasma dopamine level during training enhanced transfer as well as learning of auditory discrimination.

Approximately half of the participants (5 from the dopamine group and 4 from the placebo group) returned 20 days later for the retention test. FD performance of these participants significantly differ from the post-test (Fig. 3b; main effect of time: est. = 0.1081, s.e. = 0.044, t = 2.48, p = 0.042), but a significant group difference was still observed (Fig. 3b; main effect of group: est. = 0.4297, s.e. = 0.1109, t = 3.88, p = 0.001; group by time interaction: est. = −0.0977, s.e. = 0.0654, t = −1.49, p = 0.179), indicating successful maintenance of the dopmaine-enhanced learning. WM performance of the returning participants did not differ from the post-test (Fig. 3c, d; 2-back: main effect of time: est. = −0.1174, s.e. = 0.3041, t = 0.39, p = 0.711; group by time interaction: est. = 0.5286, s.e. = 0.4543, t = 1.16, p = 0.282; 3-back: main effect of time: est. = −0.2300, s.e. = 0.1069, t = −2.151, p = 0.068; group by time interaction: est. = −0.1761, s.e. = 0.1602, t = −1.10, p = 0.308). For Tone 3-back, a group difference was also retained (Fig. 3d; main effect of group: est. = −0.6255, s.e. = 0.2287, t = −2.74, p = 0.014), indicating successful maintenance of the transfer-enhancing effect of dopamine. Note that the retention results should be considered preliminary due to the relatively small number of returning participants.

### Working memory learning and transfer was uninfluenced by dopamine precursor during training

We next examined how WM learning was affected by elevation of plasma dopamine level during training. The dopamine and placebo groups improved similarly during Tone 2-back (Fig. 4a; training session 1-3; main effect of time: est. = 0.3396, s.e. = 0.0658, t = 5.16, p < 0.001; group by time interaction: est. = −0.0037, s.e. = 0.0901, t = −0.04, p = 0.968; main effect of group: est. = −0.0265, s.e. = 0.3232, t = −0.08, p = 0.935) and 3-back training (Fig. 4a; training session 4-9; main effect of time: est. = 0.0796, s.e. = 0.0185, t = 4.30, p < 0.001; group by time interaction: est. = 0.0211, s.e. = 0.0262, t = 0.81, p = 0.429; main effect of group: est. = −0.2610, s.e. = 0.1959, t = −1.33, p = 0.196), revealing a lack of immediate dopamine effect on learning or performance. Between the pre- and post-tests without drug administration, the two groups also improved equally for both the trained 2-back (Fig. 4c; main effect of time: est. = 1.1918, s.e. = 0.1696, t = 7.03, p < 0.001; group by time interaction: est. = −0.0950, s.e. = 0.2399, t = −0.40, p = 0.696; main effect of group: est. = 0.0036, s.e. = 0.4440, t = 0.01, p = 0.994) and 3-back conditions (Fig. 4d; main effect of time: est. = 1.1426, s.e. = 0.1365, t = 8.37, p < 0.001; group by time interaction: est. = −0.1809, s.e. = 0.1931, t = −0.937, p = 0.359; main effect of group: est. = −0.2188, s.e. = 0.3510, t = 0.62, p = 0.537), confirming the lack of dopamine effect on WM learning. Further, though WM training also improved the untrained FD task (Fig. 4b; main effect of time: est. = −0.3478, s.e. = 0.0718, t = −4.85, p < 0.001), such transfer of learning was also unaffected by dopamine manipulation (group by time interaction: est. = 0.0897, s.e. = 0.1015, t = 0.88, p = 0.387; main effect of group: est. = −0.1469, s.e. = 0.1966, t = −0.75, p = 0.459).

**Figure 4.**
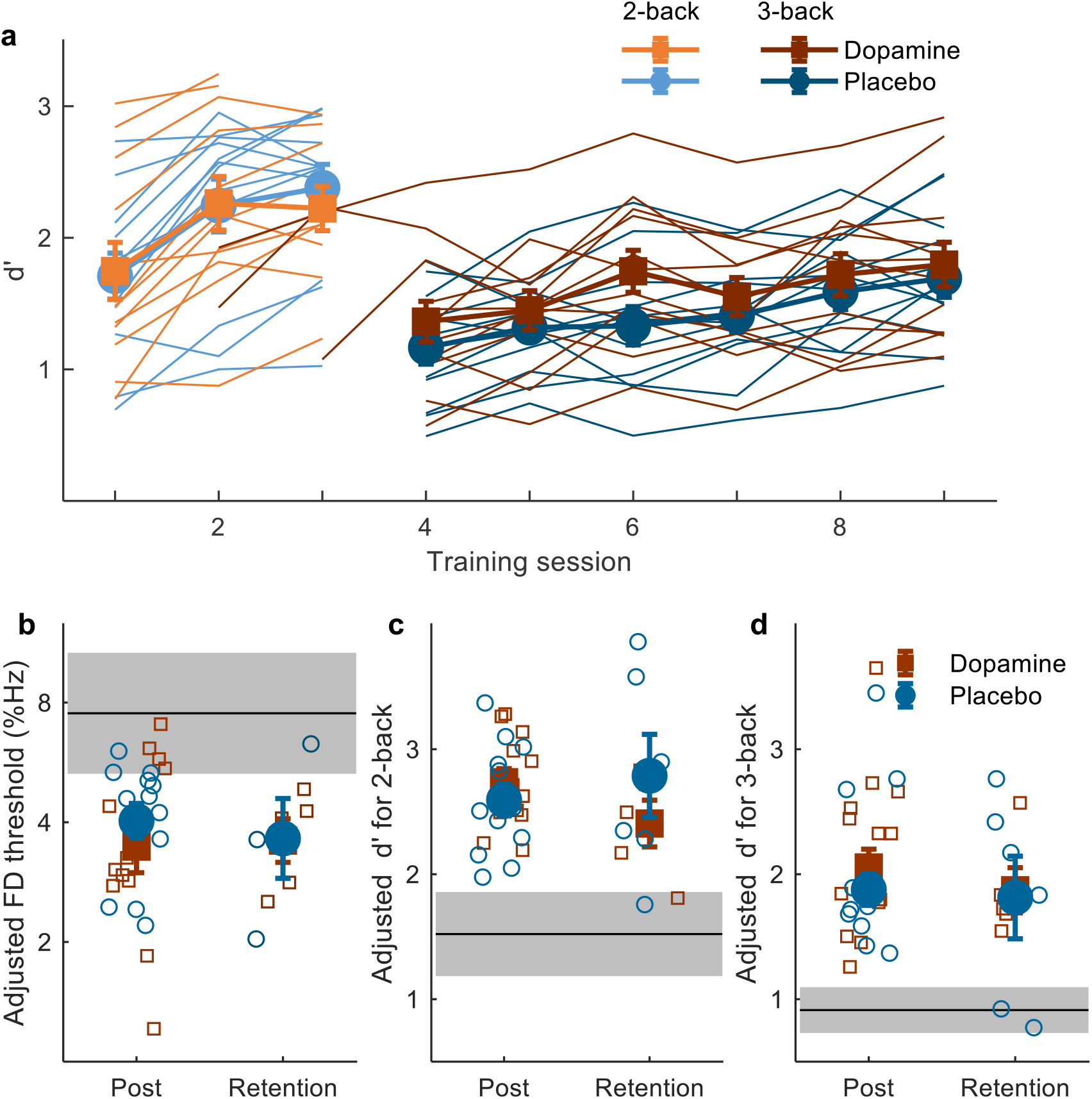
Effect of dopamine on WM learning and its transfer to FD. (a) Multiple-session training on the Tone n-back task for individuals (thin lines) and group average (thick lines with filled squares/diamonds) for the dopamine (N=12, in shades of orange) and placebo (N=12, in shades of blue) groups. n was first fixed at 2 (light colour) and then switched to 3 (dark colour) after 2-back performance exceeding 90% correct for 5 out of 8 consecutive blocks or improving for 3 consecutive sessions. (b, c, d) Individual (open symbols) and group mean (filled symbols) performance for FD (b) and WM 2-back (c) and 3-back (d) performance on post-training and retention (N=5 for dopamine group and N=6 for placebo group) tests, adjusted to account for individual differences in the pretest, with mean pretest performance (black lines) and their 95% confidence interval (grey areas) plotted within each panel. Error bars are ± 1 s.e.m.

The participants returning 20 days later (5 from the dopamine group and 6 from placebo group with 2 did not finish FD task) showed neither change of performance from the post-test or between the groups (Fig. 4b, d; Tone 3-back: main effect of time: est. = 0.0589, s.e. = 0.2290, t = 0.26, p = 0.801; main effect of group: est. = 0.0881, s.e. = 0.7449, t = .12, p = 0.907; group by time interaction: est. = −0.1156, s.e. = 0.3111, t = −0.37, p = 0.716; FD, main effect of time: est. = 0.1507, s.e. = 0.1269, t = 1.19, p = 0.256; main effect of group: est. = 0.2025, s.e. = 0.4354, t = 0.47, p = 0.649; group by time interaction: est. = −0.0850, s.e. = 0.1890, t = −0.45, p = 0.660) with the exception of 2-back performance, for which both groups had significant worse performances from the post-test to retention (Fig. 4c; main effect of time: est. = −.3803, s.e. = 0.1608, t = −2.37, p = 0.038) by similar amount (main effect of group: est. = −0.7529, s.e. = 0.5547, t = −1.36, p = 0.193; group by time interaction: est. = 0.2832, s.e. = 0.2180, t = 1.30, p = 0.221).

### FD but not WM training with dopamine affected baseline-learning and FD-WM relationships

We next examined how dopamine effect on learning varied with baseline performance, particularly to determine if the null effect of dopamine on WM training at the group level resulted from enhanced learning in some participants and reduced learning in others. For FD (Fig. 5a), the amount of learning between the pre- and posttests increased with pretest threshold in the placebo groups regardless of training task (blue circles and dashed line; main effect of pretest threshold: est. = 0.7458, s.e. = 0.1948, t = 3.83, p = 0.001; pretest threshold by training task interaction: est. = −0.1571, s.e. = 0.0890, t = −1.77, p = 0.094). This relationship was varied by dopamine during FD training (main effect of pretest threshold: est. = 1.4199, s.e. = 0.1777, t = 7.99, p < 0.001; pretest threshold by training task interaction: est. = −0.6153, s.e. = 0.0890, t = −6.9171, p < 0.001), resulting in elevated learning across the pretest performance range (red filled squares and solid line; r = 0.9117, p < 0.001), but not by dopamine during WM training (r = 0.4372, p = 0. 155). For WM (Fig. 5b), pretest performance did not affect learning in the placebo groups (main effect of pretest threshold: est. = −0.1994, s.e. = 0.6847, t = −0.29, p = 0.774; pretest threshold by training task interaction: est. = −0.1044, s.e. = 0.3687, t = −0.28, p = 0.780). Dopamine during WM training did not introduce any baseline-dependent influence (main effect of pretest threshold: est. = −0.1737, s.e. = 0.4042, t = −0.43, p = 0.673; pretest threshold by training task interaction: est. = −0.0900, s.e. = 0.2565, t = 0.35, p = 0.730), with learning amount similar to the WM placebo group at both good and poor starting performance levels. Dopamine during FD training enhanced WM in comparison to placebo, but did so independently of pretest performance (WM pretest by drug group interaction: est. = −0.2202, s.e. = 0.4164, t = −0.53, p = 0.605).

**Figure 5.**
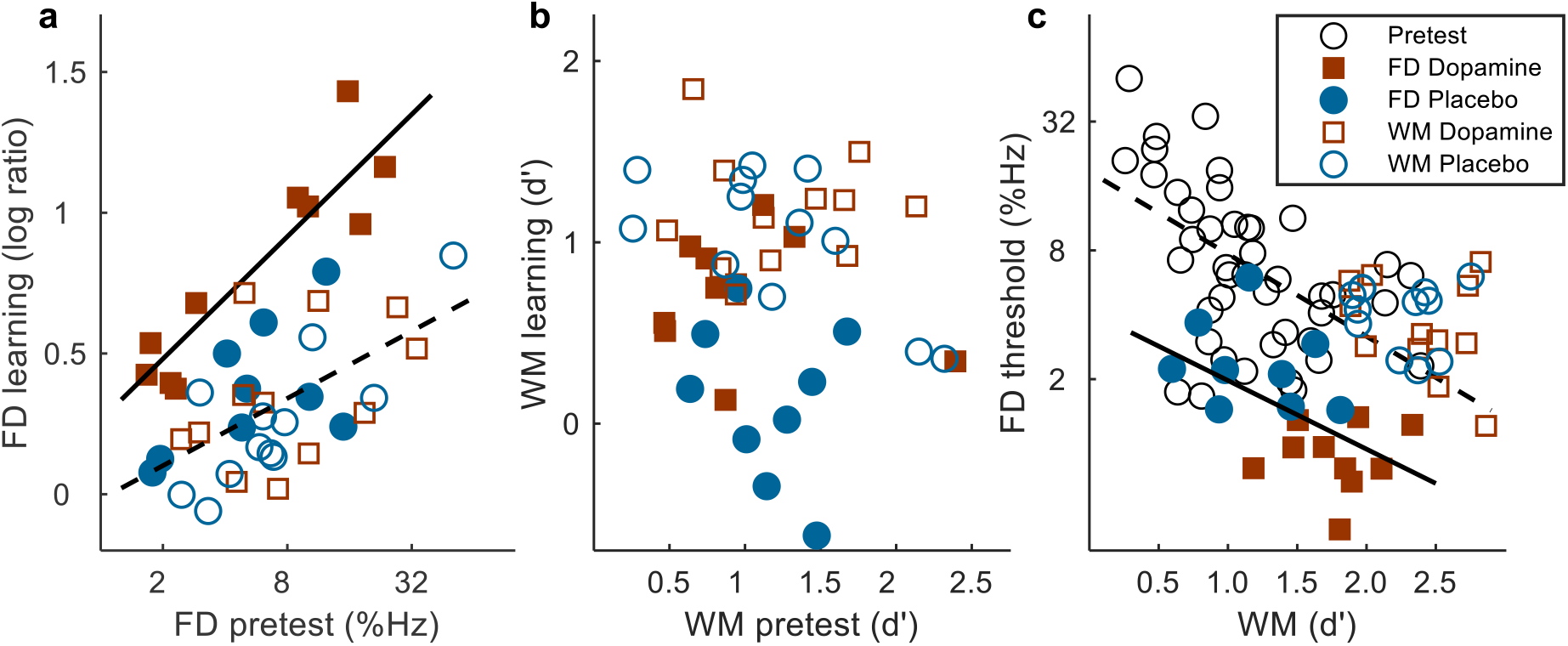
Influence of training with dopamine on baseline-learning and FD-WM relationships. Scatterplots show the relationships between pretest performance and amount of learning for FD (a) and WM (b), and the relationships between FD and WM performance (c) before and after training. FD learning (a) was calculated as log_10_(pretest/posttest). WM learning (b) was calculated as pretest-to-posttest increase in mean d’ for the tone 2- and 3-back conditions. Posttest FD and WM performance (c) was adjusted to remove individual differences in pretest performance. Significant correlations were denoted with solid and dashed lines.

Consistent with previous reports 28, FD and WM performance were correlated before training (Fig. 5c, black open circles and dashed line; r = −0.51, p< 0.001, N=43). Influence of dopamine manipulation during training on this relationship was examined using LME, with post-training performance adjusted to remove pre-training individual differences. Change of relationship between the pre- and posttests varied with training type (Fig. 5c; main effect of WM as predictor for FD: est. = −0.6453, s.e. = 0.1195, t = −5.40, p < 0.001; WM by test by training interaction: est. = 0.0887, s.e. = 0.0237, t = 3.75, p < 0.001). Follow-up analyses indicate that FD training changed the FD-WM relationship (WM by test interaction: est. = −0.2664, s.e. =0.0795, t = −3.35, p = 0.002), but WM training did not (WM by test interaction: est. = 0.0318, s.e. = 0.0546, t = 0.58, p = 0.563).

### Neural modeling of dopamine effects on auditory perceptual learning

To examine whether Madopar administered during training affected FD learning via trial-by-trial feedback, such as the reward predictor error (RPE) code carried by phasic dopamine release ^20–23^, or via long-term mechanisms such as protein synthesis dependent memory consolidation ^7,43^, a reinforcement-based reweighting neural network model was constructed for FD learning (Fig. 2; See Materials and methods for mathematical description). Such models have been recently demonstrated to be able to simulate animal as well as human perceptual learning ^18^. In particular, RPE, the proposed role of phasic dopamine release, was chosen to serve as the reinforcement signal ^38^. FD learning of the model results from changes of input weight from the sensory encoding layer to the memory layer (Figure 2). Essentially, model learning leads to activity enhancement in the memory units specific to the trained stimuli, similar to the stimulus specific recruitment of neurons beyond the primary auditory cortex after coupled electrical stimulation of midbrain dopamine neurons with pure tones observed in rat brain ^6^.

As phasic dopamine release in sensory cortex has been shown to correlate with reward, expected reward, and RPE under various situations ^2^, we simulated influence of dopamine drug on FD learning for each of these possible mechanisms (Figure 6). Each FD threshold was obtained via 50 simulations using the same adaptive track procedure as in the human training experiment. All parameters unrelated to dopamine functions were fixed after baseline FD learning was established (Figure 2B).

**Figure 6.**
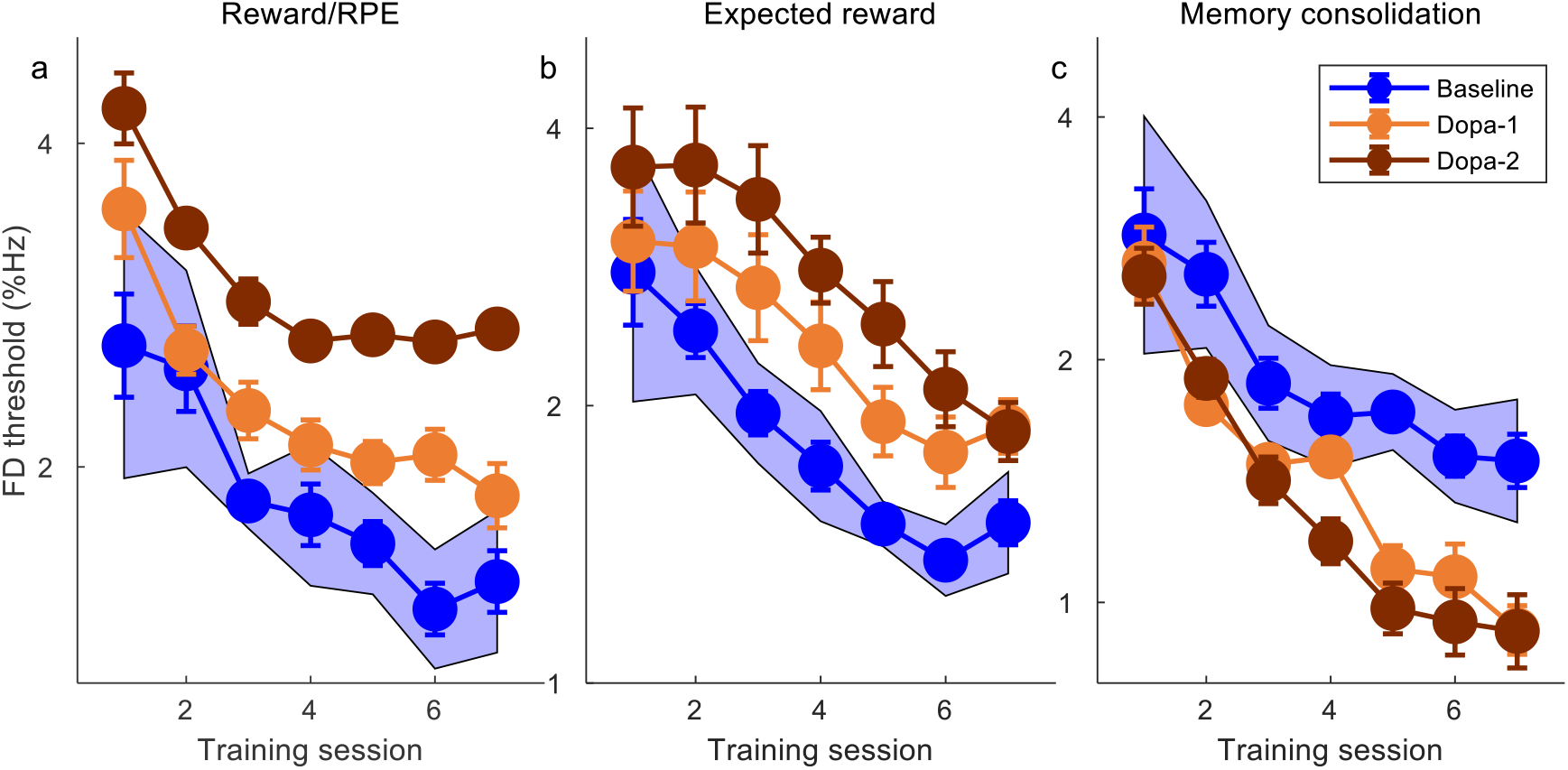
Simulated FD learning of the neural network model under different dopaminergic functions. (a) Dopamine serving as reward (R) or reward prediction error (RPE) signals after each trial. Madopar effect was simulated by a 50% (light brown) or 100% (dark brown) increase from baseline dopamine level (blue). (b) Dopamine serving as expected reward (ER) for each trial. Madopar effect was simulated by a 50% (light brown) or 100% (dark brown) increase from baseline dopamine level (blue). (c) Dopamine serving to enhance memory consolidation or reduce memory decay across training sessions. Madopar effect was simulated by reducing memory decay rate from the baseline level (blue) by 50% (light brown) and 75% (dark brown) respectively. Mean (error bars represent standard error of mean) threshold of 9 adaptive tracks in each session was obtained from 50 simulations for each condition. For baseline conditions, shaded areas represent 95% confidence intervals of learning curves. All parameters unrelated to dopaminergic functions were fixed in all simulations.

Because RPE was calculated as difference between actual reward and expected reward (equation 5), an additive increase in actual reward was equivalent to the same additive increase in RPE. Therefore, reward (R) and RPE mechanisms were simulated together. The amount of drug-induced change in dopamine level was simulated to be 0-200% of steady dopamine level before drug administration in the step of 25%. When dopamine served as reward or RPE signal for learning (Figure 6A; for clarity, only baseline, 50% and 100% increases were plotted), dopamine drug administration during training led to discrimination threshold elevation (performance impairment) and earlier performance plateau. The drug effect increased monotonically with increasing magnitude of drug-induced increase in dopamine level. When dopamine served as signal for expected reward (Figure 6B), dopamine drug also elevated FD threshold, but with little influence on learning magnitude. In contrast, when dopamine served to enhance learning consolidation across sessions, dopamine drug led to no performance change at the beginning of training, but enhanced learning particularly during later sessions (Figure 6C). Thus, function of dopamine drug via reward/RPE, expected reward, or consolidation manifested distinctive patterns of influence on FD learning, among which only the pattern for the consolidation mechanism matched that of human behavior (Figure 3).

## Discussion

We combined psychopharmacological experiments and neural network modelling to examine how dopamine drug influences multi-session perceptual learning in healthy human adults. In two parallel double-blinded experiments, a dopamine precursor drug, Madopar, was administered before each of multiple daily sessions training either auditory perception or working memory (WM). Compared to placebo, Madopar enhanced auditory perceptual learning of a tone frequency discrimination (FD) task and its transfer to WM even when tested outside the time window of plasma dopamine elevation, which appears retained for at least 20 days. In contrast, the same dopaminergic manipulation during WM training had no effect on learning or transfer. A neural network model of FD learning revealed three distinctive behavioural modulation patterns for three proposed dopaminergic functions in the auditory cortex: reward /reward prediction error (RPE), expected reward, and memory consolidation. Only memory consolidation provided a match between model simulations and experimental observations, indicating that dopaminergic modulation of multi-session perceptual learning may be mediated by enhancement of memory consolidation across sessions.

### Dopaminergic modulation of human learning and cortical plasticity

The finding that Madopar enhanced auditory perceptual learning and transfer is consistent with and extends previous reports of modulation of human learning by dopaminergic drugs in healthy adults or patients of Parkinson Disease, including reward learning ^27,44–46^, motor learning ^13,14,47^, and word learning ^15^. Thus, drug-induced changes in plasma dopamine level, lasting from minutes to hours, can enhance learning across different tasks, modalities, and intrinsic tonic dopamine levels (reduced in patients with Parkinson Disease in comparison with healthy adults), suggesting a widely present mechanism in the brain for slow, non-phasic dopamine releases to affect neural plasticity.

The lack of dopaminergic effect on WM learning, however, presents an exception to this literature. Note that variation of dopaminergic effect is associated with training experiences, but not with task type. As a matter of fact, WM was enhanced by Madopar administration during FD training, as transfer from the enhanced auditory perceptual learning, but not by Madopar administered during WM training. This distinction is important because auditory perception and working memory appear intertwined not only at the behavioural level, as evidenced by performance correlation and bi-directional transfer of learning between FD and Tone n-back tasks observed here and previously ^28^, but also at the neurophysiological level, in that the auditory cortex subserves perceptual as well as memory functions ^48–51^. In terms of training-induced learning, accumulating evidence points to distinct neural substrates: the striatal-prefrontal dopaminergic network for working memory ^30^ and the auditory cortex for frequency discrimination ^52–54^. Under this light, the contrasting effects of Madopar in the two training experiments suggest that dopamine serves a plasticity-modulating function in auditory cortex, but only for training or experience dependent plasticity. While training auditory working memory certainly engages auditory cortex as well, training-induced neural modifications would take place elsewhere, presumably in the striatal-prefrontal dopaminergic network, leaving auditory cortex irresponsive to the plasticity-modulating effect of dopamine.

Modulation of auditory cortical plasticity by dopamine is consistent with existing evidence. A number of animal studies have shown that dopamine can trigger tonotopic map plasticity in auditory cortex during associative learning ^2,10,55^, to the effect that cortical representation area of sounds was adjusted in response to their ability to predict dopamine release. In humans, levodopa was reported to enhance auditory cortical activity during reward learning without differentiating rewarded and unrewarded stimuli ^17^. While the exact mechanism of dopaminergic modulation of sensory plasticity remains under debate, there are evidence for multiple, non-exclusive roles of dopamine in sensory cortices, including reward related learning signals and memory of behavioural significance ^2,56^. Specifically, for multi-session auditory perceptual learning, animal studies suggested that dopamine in the auditory cortex modulates learning via protein synthesis dependent processes of memory consolidation ^7,43^. The neural network model, designed to distinguish contributions of these mechanisms, support that in humans, the observed modulation of perceptual learning by dopamine drugs is likely mediated by enhancement of memory consolidation across training sessions. Though other dopaminergic functions may be at work simultaneously, the current study provides, to our knowledge, the first evidence for long-term dopaminergic modulation of memory consolidation in humans.

### Implications for the role of dopamine in learning theories

Computationally, learning can happen in two distinct forms: ‘model-free’ learning via instantly reinforced stimulus-response associations, and ‘model-based’ learning that takes into account knowledge, or “model” of the environment and potential action outcomes in long term ^57^. While it is well established that the sub-second phasic dopamine releases serve as reinforcement signal for model-free learning by coding reward prediction error ^58^, relatively little is known about the role of dopamine in model-based learning. Employing paradigms designed to dissociate model-free and model-based behaviours, dopaminergic drugs have been shown to promote model-based over model-free choice in healthy, young adults ^46^, and to remedy impairment of model-based learning in patients with Parkinson’s disease off medication ^27^. The dopaminergic effect on model-based learning was suggested to be mediated by WM by Sharp and colleagues ^27^ based on the observation that model-based learning was correlated with WM when the patients were on dopamine medication, but not when they were off medication. Indeed, auditory WM and perception are correlated, and their learning transfers bi-directionally, in healthy adults ^28,29^, consistent with neurophysiological observations of engagement of auditory cortex for both tasks ^48–51^. The current results further revealed that the relationship between WM and auditory discrimination changed after FD, but not after WM training (Fig. 4), suggesting that the shared neural processes giving rise to performance correlation coincide with the FD learning substrate. However, neither performance nor learning of WM was affected by the same dopamine manipulation that enhanced FD learning, demonstrating dissociation of dopaminergic effect on their respective learning mechanisms despite the intertwined relation of the two tasks.

As an attempt to disentangle the multiple possible roles of dopamine in learning, a neural network model incorporating both model-free and model-based learning processes was constructed for FD learning (Figure 2). Model-free learning occurred across trials within each training session, with reward prediction error (RPE) served as trial-by-trial reinforcement signal to guide synaptic weight changes from a tonotopically organized encoding layer to a memory (delayed response) layer. Performance changes between sessions reflected model-based learning, in that previous experiences (long-term memory) were retained (consolidated) to certain extent and would impact subsequent learning. Model simulations identified distinctive learning modulation patterns by dopamine drug for dopamine serving as reward/RPE, expected reward, and memory consolidation modulator, only the last matching that observed in training experiments. Thus, though dopamine is capable of modulating both model-free and model-based learning via different mechanisms, its effect on multi-session perceptual learning in humans is most likely mediated by long-term memory consolidation.

The combined results of neural modelling and behavioural training thus point to dopaminergic modulation of long-term memory consolidation in sensory cortices as underlying learning and transfer enhancement by dopamine precursor drugs. In doing so, the results support a unified learning modulatory role of dopamine across time scales: sub-second level fluctuations may serve as learning reinforcement signal for model-free learning by trial and error, while longer-term (minutes to hours) changes modulate the influence of previous experiences, so that behavioral adaptations reflect both current environmental demand and previously accumulated knowledge.

### Limitations

Out results have limitations that could be addressed in future work. Participants in our study were predominantly female (only 4 males in total). Previous studies in humans have revealed gender differences in striatal dopamine release ^59^ and extrastriatal dopamine D2-like receptor expressions ^60^. Given the uneven gender distribution of the volunteer participants, it is unclear whether our results can be generalized across genders. Further work with more gender-balanced sample could clarify this point. In addition, the relationship between dopamine level and behavioural performance has been suggested to follow an “inverted-U shape”^24^, where a dopaminergic drug may cause beneficial or detrimental effects depending on baseline dopamine levels of individuals ^61,62^. The current experiments employed a single level of Madopar dose that has been suggested to enhance learning behavior by the existing literature, leaving dose effect on learning unexamined. Also, the plasma dopamine level of participants was not directly monitored through training. Further studies with multiple dose levels and direct monitoring of plasma dopamine level would be needed before dopamine drugs can be used to enhance learning in clinical or educational settings.

## Acknowledgments

We thank Ting Huang and Ting-Ting Yan for assistance with data collection, and an anonymous reviewer for helpful comments on data analyses.

